# Multisensory perceptual learning in virtual reality may facilitate transfer to untrained locations but is impacted by training procedures

**DOI:** 10.1101/2023.05.04.539103

**Authors:** Catherine A. Fromm, Krystel R. Huxlin, Ross K. Maddox, Melissa J. Polonenko, Gabriel J. Diaz

## Abstract

Visual training improves performance in visually-intact and visually-impaired participants, making it useful as a rehabilitation tool. An interesting question in rehabilitation is whether invocation of multisensory integration could increase training efficacy. To investigate, participants completed a 10-day training experiment wherein they repeatedly performed a 4-way global motion direction discrimination task. On each trial, participants were presented with a 5° diameter visual global motion stimulus placed in a gaze-contingent manner at 10° azimuth/elevation. The visual-only group was presented the unimodal visual stimulus. However, for the auditory/visual (AV) group, the visual stimulus was paired with a pulsed white-noise auditory cue moving along in a direction consistent with the horizontal component of the visual motion stimulus. Direction range thresholds (DRT) were computed daily. The motion direction discrimination learning transferred fully in both groups, regardless of the presence of a feature-based attention cue. However, the figure-ground segregation learning only fully transferred to untrained locations with the addition of the feature-based attention cue. These results speak to the multiple levels of processing on which perceptual learning can operate.

**PACS:** 0000, 1111

**2000 MSC:** 0000, 1111

## 1. Introduction

As an individual grows, their visual system must develop in response to ever-changing demands. This does not stop when adulthood is reached, and perceptual ability can change throughout a lifetime. Visual perception requires adaptation to different situations and optimization of processing to better fit the statistics of the natural visual environment. These changes can be short-term adaptations, which quickly fade when conditions change, or can take the form of longer-lasting changes due to plasticity. Plasticity in the brain is essential to the growth and development of cognition and can be accessed through a process referred to a *perceptual learning*, in which the learner’s actual perceptual ability is altered through specific training. Perceptual learning has also been demonstrated in laboratory settings in response to specific visual training regimes, and has been used as a method to rehabilitate visual deficits, particularly in hemianopia (Huxlin, 2008; Sahraie et al., 2006; Elshout et al., 2016; Bergsma et al., 2012).

There is interest in using perceptual learning as a form of non-invasive rehabilitation for certain forms of vision loss. For example, hemianopia is a form of cortical vision loss often caused by stroke in the primary visual cortex (Goodwin, 2014). It manifests as the loss of conscious vision in half of the visual field contralateral to the damage. This damage can be partially rehabilitated through perceptual training at targeted locations at the edge of the intact visual field (Melnick et al., 2016; Saionz et al., 2022, 2021). However, In certain training contexts, the learned benefit is specific to the trained stimulus dimension like orientation (Ahissar and Hochstein, 1997), or retinotopic location (Karni and Sagi, 1991). This is thought to result from improvements only in populations of neurons at the earliest levels in processing which have specificity for certain stimulus features or spatial locations (Sagi and Tanne, 1994). Particularly in a rehabilitation context, the specificity of training can make the process of re-learning to see slower and less efficient, as each individual region in the visual field as well as each visual task must be trained in series.

There has been some success mitigating the spatial specificity of perceptual learning through the use of alternative learning paradigms which may shift the mechanism of learning to a higher level in the processing hierarchy (Dosher et al., 2013). These methods include double training, where a separate irrelevant task is trained simultaneously in a second location, (Wang et al., 2014), the invocation of visual attention (Donovan et al., 2020; Hung and Carrasco, 2021) and other strategies designed to shift processing away from the spatially specific primary visual areas. Similar to visual-only perceptual learning, multisensory perceptual learning is also possible. After repeated exposure to multi-modal stimulus, perceptual performance on an audiovisual task improves (Shams and Seitz, 2008). Multisensory audiovisual training has also been shown to generalize to visual improvements, an indication that mechanisms where multisensory integration occurs may be implicated in the learning (Seitz et al., 2006; Kim et al., 2008; Powers III et al., 2016a). Training with audiovisual stimuli may be a good candidate to improve learning efficiency especially when the visual modality is compromised.

The neurological mechanisms of audiovisual learning seem to extend beyond modulation of primary sensory cortices alone (Bavelier and Neville, 2002). Training with auditory-visual stimuli produced enhancement in the superior colliculus, but only when the auditory and visual cues were presented in a temporally synchronous and spatially congruent manner necessary for audiovisual binding. (Grasso et al., 2016; Noel et al., 2018). The superior colliculus has also been implicated in spatial coordination as well as audiovisual processing (Lee and Groh, 2014; Groh et al., 2001). In addition to the superior colliculus, area MT has been implicated in both motion processing and audiovisual integration (Kolster et al., 2010), and has shown potential as a mechanism in multisensory training (Grasso et al., 2016)

The present study examines the potential benefits of multisensory integration for learning in intact individuals. Specifically, it has been designed to test whether audio-visual training enhances visual motion direction discrimination learning, whether this learning is retained when the audio cues are removed, and whether this learning transfers across the visual field, benefiting untrained visual field locations.

## 2. Methods

### 2.1. Participants

Participants were recruited from the Rochester Institute of Technology (RIT) campus community. The experiment was approved by the Institutional Review Board at RIT and all participants gave informed consent prior to participation. Participants had normal or corrected-to-normal vision, and self-reported as having no documented hearing or auditory processing disorders. A total of 17 participants, 6 females, participated in the full 12 days of the study. The mean age (±SD) was 25.2 ±3.2 years. Six participants (3 females) attended the first session only and opted to self-exclude due to discomfort using the virtual reality system or were excluded by the experimenter because of unusually poor task performance in the practice block.

### 2.2. Apparatus

During all experiment sessions, participants were seated wearing an HTC Vive Pro Eye virtual reality headset. The binocular field of view of the headset is nominally 110° horizontally, but this varies with the adjustable distance of the eyes from the headset’s optics. During pilot testing, we verified that the task environment was able to maintain the nominal update rate of approximately 90 Hz. The experimental software was created and rendered using Unity3D version 2019.1.14f1, and run on a PC equipped with an Intel i7-6700 CPU and an NVIDIA 2080 RTX graphics card. The eye tracker integrated into the HTC Vive Pro Eye was used during each session, and controlled using the manufacturer-provided SRAnipal plugin to Unity3D, version 1.3.2.0. Auditory stimuli were created using the SteamAudio audio spatializer plugin and spatialization was conducted using a generic head rotation transfer function (HRTF). The HRTF in this context is the function used to compute the spatial effects of the human ear on the virtual sound waves so that virtual audio presented with headphones sounds as if it is truly coming from a distant point in space. A generic HRTF was sufficient for our purposes because participants were only asked to make coarse judgments about the auditory portion of the task, as discussed in Section 2.5.

At the start of each testing or training session, participants adjusted the headset to improve comfort and display quality, and then completed the integrated Vive Pro Eye calibration routine. Following calibration, a custom calibration assessment was run through Unity. Participants were sequentially presented with each target of a grid of nine spherical targets that were equally spaced across a visual angle of 5°. Targets were yoked to the head position. All targets were at a virtual depth of 0.57m, which was experimentally determined to be the virtual image plane of the optics in the specific headset used for this experiment. This depth was selected to minimize the effects of the vergence accommodation conflict which has been known to cause discomfort during use of virtual reality displays (Hoffman et al., 2008). Data was recorded for a period of 500ms during fixation at each target, and error was measured between the true target direction and the gaze direction reported by the eye tracker. The average error across all sessions and all participants at the central fixation point was 1.1° with a standard deviation of 0.4° of visual angle.

Once the calibration assessment concluded, the seated participant was instructed to adopt a comfortable head pose that could be maintained for a long period of time. Head position was to remain constant during the trial to maintain consistency in the auditory stimulus localization cues. The participant’s head position in the 3D world reference frame was recorded in Unity3D, and used to constrain the initial head position to this pose at the start of each trial, similar in spirit to the use of a fixation point to constrain gaze direction. If the head position rotated from it’s original orientation by more than 2 °, a realignment procedure began. To realign the head, participants moved their head to align a rectangular box that was stationary in head-centered reference frame with a bar that was stationary at the world reference frame. This world-fixed box was located at the fixation point, parallel to the ground, and perpendicular to the vector between the head and the fixation point. Trials would not begin unless fixation was within 0.3° of the fixation point, and head pose was within 2° of the world-fixed box for one uninterrupted second.

Fixation and head pose were also constrained during stimulus presentation. If the head position deviated from the home position by more than 2° of rotation in any direction or translation by more than 10 cm of translation while a trial was in progress, the trial would be aborted and the stimulus reset with the same conditions. If gaze location as reported by the eye tracker deviated by more than 1.5° from the fixation point, the trial was aborted and the stimulus reset.

### 2.3. Task

Participants were asked to discriminate the global motion direction of a field of dots moving within a 5° diameter aperture. The dots could move to one of four oblique directions: upper right, lower right, upper left, or lower left. The individual dot motion directions were perturbed away from the oblique direction described in Section 2.4 to increase the direction range of the global motion over 10 discrete levels. This direction range level changed each trial following a 2-up-1-down staircase procedure. One group received visual-only stimuli for the duration of the training period, while the other group had an auditory stimulus paired with the visual dot motion during training. The auditory stimulus, described in Section 2.5, always moved with the same direction as the horizontal component of the visual dot motion as shown in Figure 1. For example, if the visual motion was directed to the upper left oblique, the auditory stimulus would move from right to left along the horizontal axis. This ensured that the group with audiovisual training stimuli had partial information about the visual motion direction.

**Figure 1:**
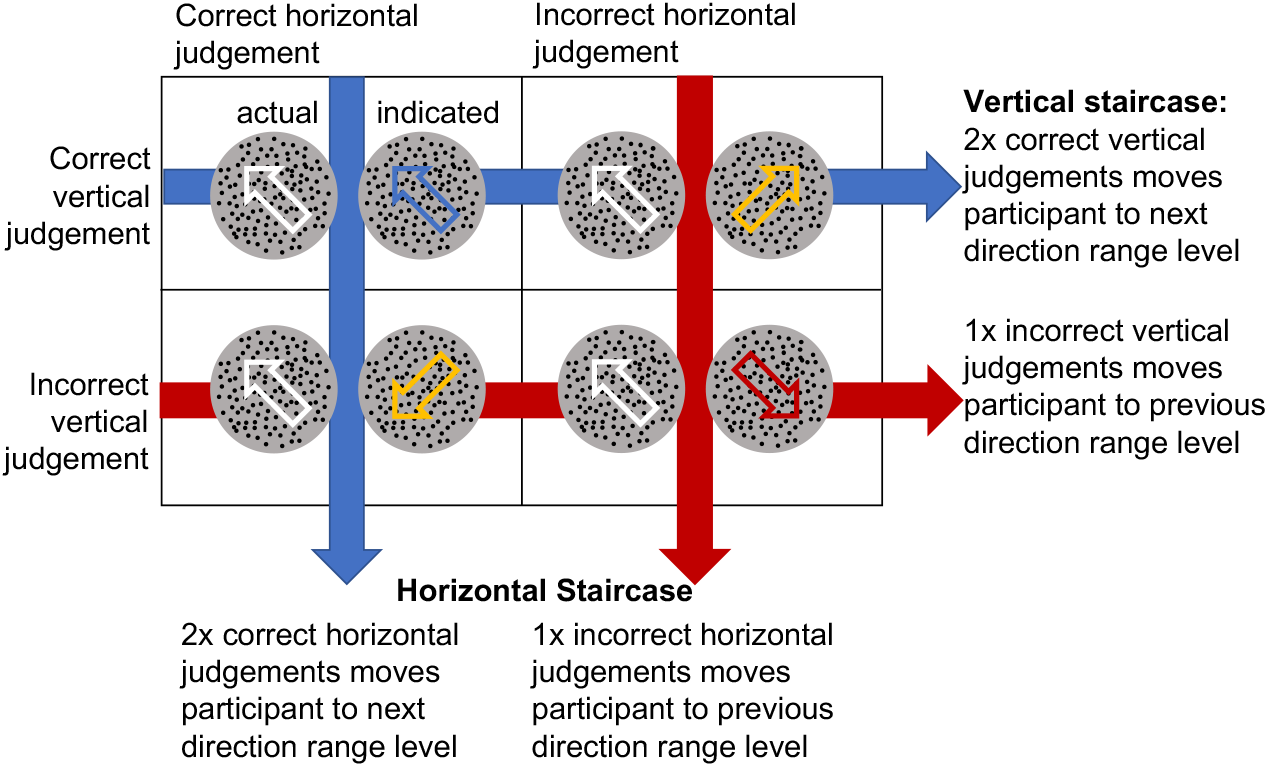
Decomposition of horizontal and vertical components of global motion stimulus. The global motion is directed to one of the four corners overall, but participants are scored separately on the horizontal and vertical components of their response.

### 2.4. Visual stimuli

We used random dot stimuli within a 5° diameter aperture. Individual dots were 14 arcmin in diameter and dot density was 3.5 dots/°^2^ . Dots moved at 10°, and had a lifetime of 250ms before disappearing and re-spawning in a new random location. Total stimulus duration was 500 ms. Difficulty of the visual task was parameterized in two ways. The first method was to manipulate the proportion of “noise” dots, which moved with random trajectories as opposed to the “signal” dots, which moved with coherent trajectories. The proportion of random dots in the stimulus was determined and set at the beginning of the experiment (in the titration block) for each participant. Once set, it remained unchanged throughout the 12-day experiment. This proportion was set according to a pre-experiment titration procedure for the purpose of modifying the baseline noise dot level of the task in order to account for individual differences. This procedure is described in Section 2.6. The second stimulus manipulation consisted of varying the range of directions in which signal dots could move. This was dynamically changed within a session, using a staircase procedure tied to performance. The direction range of a stimulus described the bounds on a uniform distribution of angles from which signal dot trajectories could be selected. For example, for a stimulus with direction range of 80°, the motion direction of a given signal dot was chosen from a distribution that spanned ±40° about one of the four oblique base directions of the stimulus (upper right, lower right, upper left, lower left).

### 2.5. Auditory Stimuli

For participants in the audio-visual training group, auditory stimulation was added during visual stimulus presentation. This auditory stimulus provided information about only the horizontal motion component of the visual stimulus on any given trial. The spatialized auditory stimulus was implemented with the SteamAudio spatializer plugin for Unity3D. The virtual sound source was defined as point source with a radius of 2.5°. The motion of the source was from a point 20° left of the visual stimulus center, to a point 20° right of center, through an arc with radius 0.57m parallel to the horizontal ground plane, as shown in Figure 2. The speed of motion of the audio source was 80°/s. The motion direction of the auditory stimulus always matched the horizontal motion direction component of the visual stimulus. For example, if the movement direction of the visual stimulus was towards either the top-right or the bottom-right, the auditory stimulus would sweep from left to right. The participants in the audio-visual training group were informed that the auditory stimulus would always match the horizontal component of motion of the visual stimulus.

**Figure 2:**
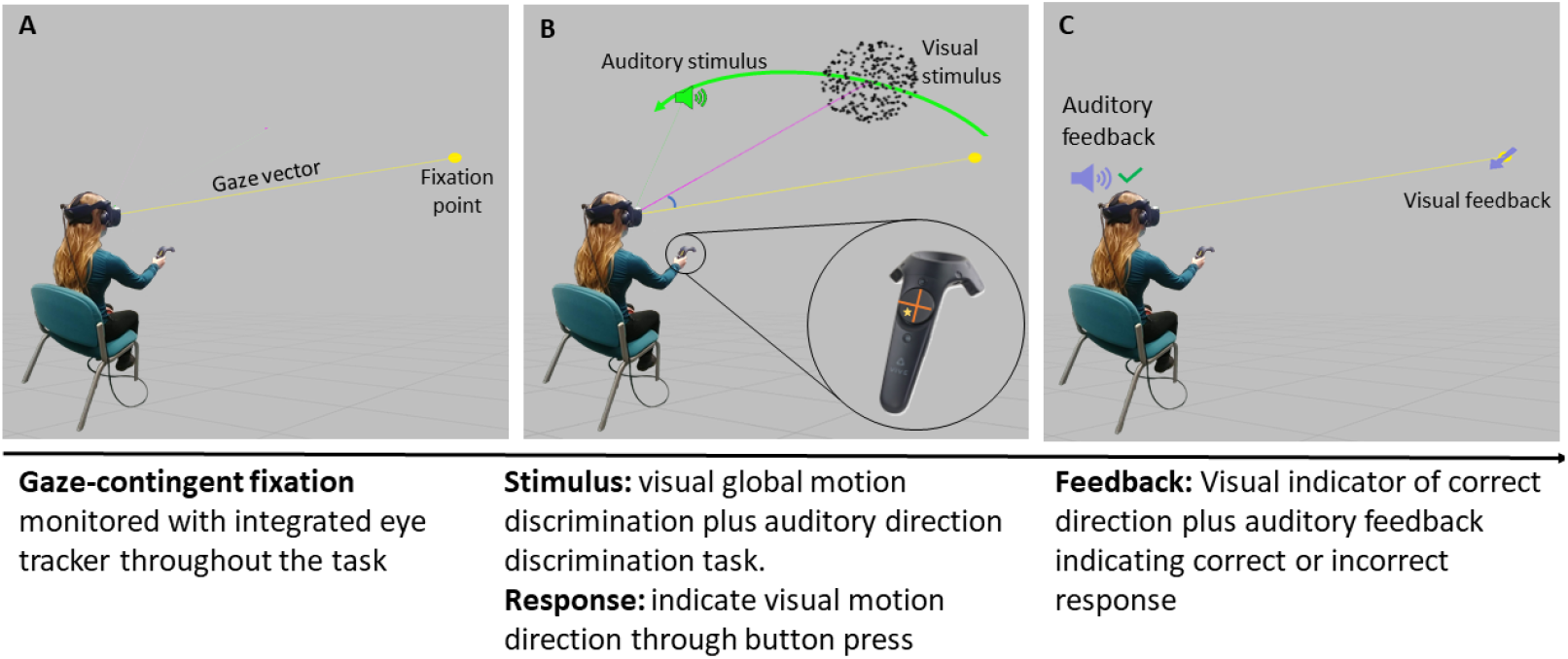
The Experimental Task: Participants were seated and wearing the HTC Vive Pro Eye virtual reality display. All stimuli were gaze contingent, and fixation was enforced with the built-in eye tracker. They viewed a visual stimulus and heard the auditory stimulus in the 3D environment, and responded with a button press

The auditory stimulus was parameterized such that perception of its motion direction would be significantly above threshold and that all participants would be able to clearly discern the cued direction. The sound used was a pulsed white noise with a duration of 500ms and pulse frequency of 12Hz. As discussed in Section 2.2, a generic HRTF was used for this experiment. As computing individualized HRTFs is an arduous process, and the benefits of using them are primarily seen when making absolute spatial localization judgments (Berger et al., 2018), we found that using the generic HRTF did not prevent participants from being able to accurately and consistently judge motion direction of the auditory source. Further, the decision to restrict the movement of the auditory stimulus to the horizontal direction rather than vertical was motivated, in part, by the finding that there is potential for up/down confusion when using generic HRTFs in virtual spatialized audio (Wenzel et al., 1993; Berger et al., 2018) but more clarity in left/right discrimination.

### 2.6. Procedure

#### 2.6.1. Setting the initial noise level for the visual stimulus

The first day of the experiment involved a procedure for the purpose of modifying the baseline percentage of random noise dots of the task to ensure that all participants started with similar performance at the outset of the study. Participants first completed 20 trials of practice with initial difficulty was set to 30% of the dots to move with completely random trajectories to familiarize themselves with the apparatus and task (see Section 2.4 for further details on random dot settings). If the practice block was so difficult that the participants could not progress, or so easy that every trial was correct, the proportion of random dots was adjusted in increments of 10% random dots until the participant could correctly respond to trials at 80° of direction range within the first 20 trials. Following this coarse adjustment of the baseline random dot ratio, participants performed a titration block consisting of 100 trials. Performance in this block was used to fine tune the random dot ratio for the training and testing blocks. After the titration block was completed, participants took a brief break out of the headset while the experimenter fit a psychometric function to their trial-by-trial results (see Section 2.8 for further details). If the threshold of the psychometric function was between 100° and 200° of direction range, this difficulty setting was kept for the remainder of the study. If it was above or below, difficulty was adjusted by adding or subtracting 5% from the random dot proportion and the experiment proceeded.

### 2.6.2. Pre- and post-tests

After the titration block, participants completed a pre-test consisting of 200 4AFC direction discrimination trials at both the training location (10° to the right and 10° up from fixation and 0.57m from the head) and at a second untrained location, 10° left and 10° up from fixation, at a depth of 0.57m (Figure 3, Day 1). On the subsequent 10 days of training (Figure 3, Days 2-11), participants completed 300 training trials in their assigned condition (visual-only or audio-visual). On the final day, (Figure 3, Day 12) participants performed post-tests in the trained and untrained locations that were identical to baseline, pre-test on Day 1. During pre- and post-tests, trial direction range was determined on the fly based on two interleaved staircases, each with a 2:1 ratio. One staircase tracked correctness of the judgment of horizontal component of the visual motion stimulus while the other tracked correctness of the vertical component judgment. When participants scored 2 correct trials in a row at a particular direction range level on either the horizontal or vertical components, depending on which staircase that trial corresponded to, direction range in the stimulus was increase by 40°, making the task harder. When they scored an incorrect response, direction range was decreased by 40°, making the task easier.

**Figure 3:**
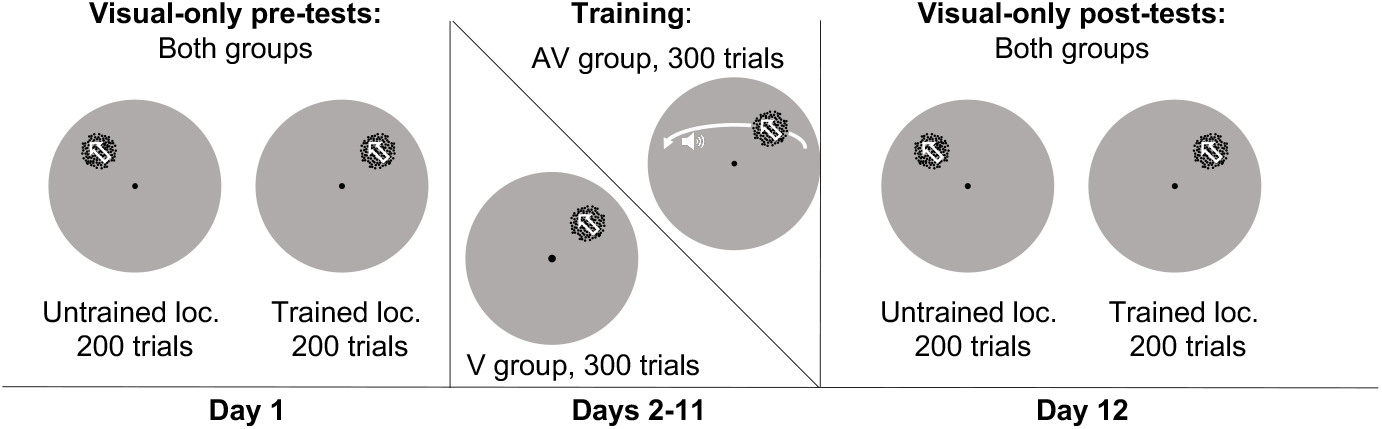
Study design: In both the visual and the audiovisual groups, the first day of the study was a single day with two blocks of 200 trials each. The first block used a training location at 10° azimuth and 10° elevation relative to the fixation point, and the second block was at -10° azimuth and 10° elevation. For the following 10 days of training, the both groups completed 300 trials per day in a single block at [10°, 10°] and the audiovisual group had an added sound cue on each trial. On the final day, both groups repeated two blocks of 200 trials each at [10°, 10°] and [-10°, 10°].

**Figure 4:**
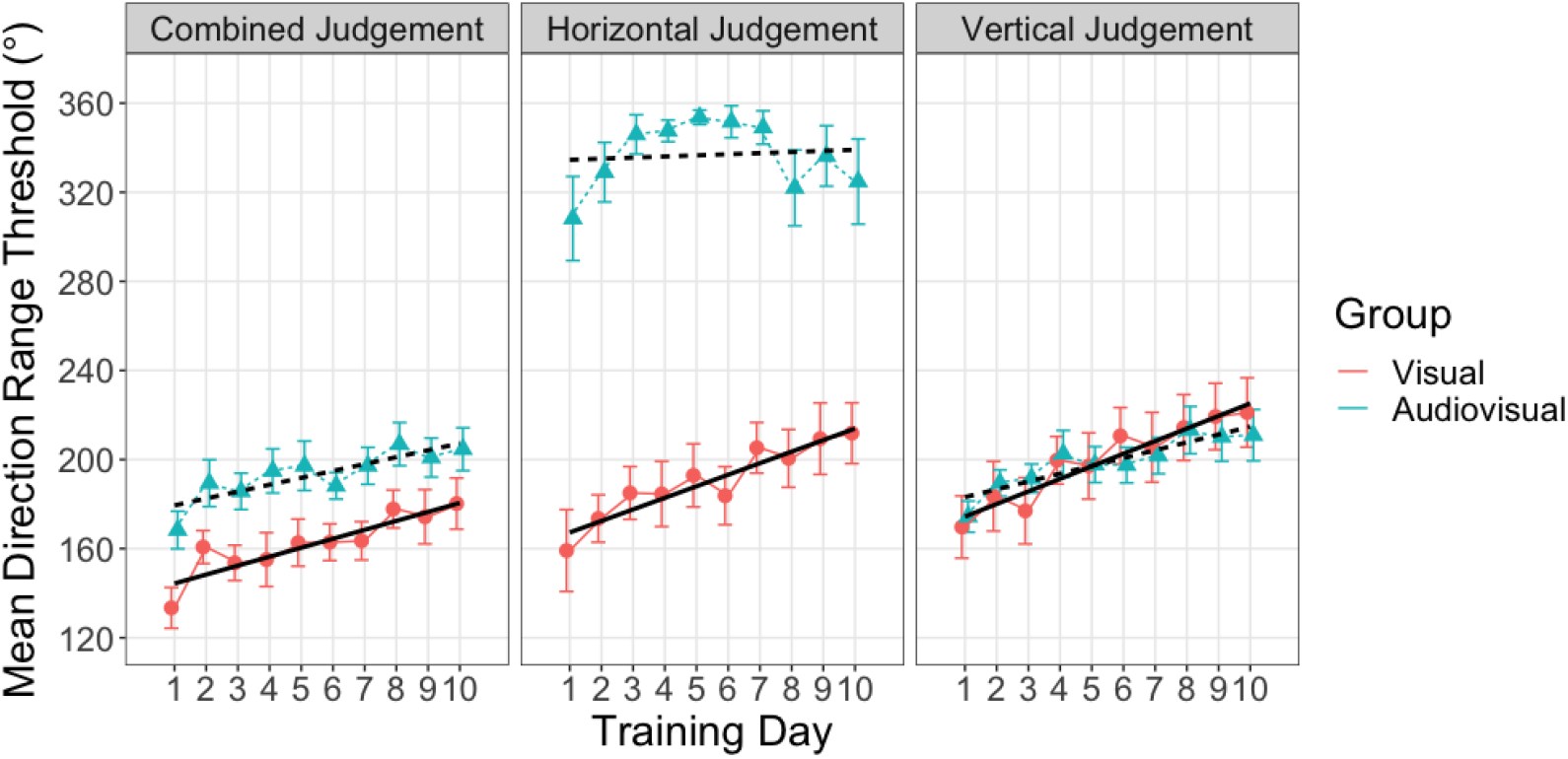
Training results: Direction range thresholds for the overall combined motion direction judgment as well as the horizontal and vertical components of the motion direction judgment are averaged for each group (error bars are standard deviation) and shown for each of the 10 days of training. The audiovisual group had n = 9 participants, and the visual-only group had n = 8 participants. Horizontal judgments for the AV group (triangle symbols, center panel) are at ceiling due to the presence of the informative auditory cue.

### 2.6.3. Training sessions

Each training session (Days 2-11) consisted of a single block of 300 trials of 4AFC direction discrimination. At the start of each training trial, a fixation point was presented in the center of the participant’s visual field at a virtual depth of 0.57m. After the eye tracker recorded 1s of fixation on the target, and the head tracker simultaneously recorded 1s of stationary head in the home position, the trial proceeded to the stimulus. If the eyes or head deviated during this 1s interval, the trial reset. Additionally, if the eyes or head deviated from their positions at any time during the stimulus presentation, or before the participant provided a response, the trial reset. After the 1s interval had ended, during continued fixation, a visual or visual+auditory motion stimulus was presented for 500ms at the same depth as the fixation target, 10° azimuth and 10° elevation away from fixation (trained location, Figure 2A). The participant was able to indicate perceived motion direction (upper right, lower right, upper left, lower left) at any point following stimulus presentation by selecting one of 4 corner regions on the HTC Vive controller’s trackpad. There was no limit placed on the response time. After the response was given, participants received feedback in the form of a blue arrow presented at fixation indicating the correct direction of the motion, as seen in Figure 2. They also received positive or a negative auditory feedback. This feedback was delivered in the form of sounds similar to those used in video games, and easily discerned as indicating positive or negative feedback.

### 2.7. Feedback

In this task, feedback was provided in multiple ways, both through an indicator sound which was positive for a correct vertical judgment and negative for an incorrect vertical judgment, and by a visual representation of the correct motion direction as an arrow. However, during training, participants were also informed of the nature of the staircase, and knew that if they got the vertical component of the task on three consecutive trials, they would move to a higher direction range trial. Before training, participants were instructed that their goal was to reach the highest direction range level possible.

Though the audible and visual feedback remained the same in the pre- and post-test sessions, the staircase procedure was changed. In the test sessions, participants were tested with two interleaved staircases, which switched randomly from trial to trial. They were informed that the level of the trials would generally improve the more overall correct responses they gave, and that both the horizontal and vertical components were important to progress, but that direction range would be somewhat randomized. This was done to ensure participants did not discount the importance of one component during testing, but also means that the task was implicitly and subtly different during testing due to slightly different feedback conditions.

### 2.8. Analysis

To assess performance in each session, data was exported from Unity3D as a custom CSV file then imported into Matlab version R2018a for quantitative analysis. Psychometric functions were fit to each session using custom code to calculate the session’s direction range threshold (DRT). DRTs were calculated by fitting a Weibull function to the percentage of correct judgments made at each level of direction range.

Weibull function parameters were fit using maximum likelihood estimation implemented in MATLAB using the Optimization Toolbox. The chance level for each component judgment was 50% and the fits were performed assuming a 5% lapse rate. The Weibull function was evaluated to find the direction range level at which participants answered correctly 72.5% of the time, the value halfway between ideal performance considering the lapse rate and chance.

Although participants made a judgment on both vertical and horizontal components of the global motion stimulus at once by selecting one of four oblique motion directions, separate thresholds were calculated for judgments of the vertical component direction and judgments of the horizontal component of motion for each training session. For example, if the correct direction was to the upper right, and the participant selected the upper left, the participant would score correctly on the judgment of the vertical component, but incorrectly on the judgment of the horizontal component (see Figure 1).

Vertical and horizontal training rates were calculated as the slope of a linear fit to the mean DRT across the 10 training sessions. The decision to analyze vertical motion direction threshold separately from the horizontal threshold made comparison of training efficacy between the unisensory and multisensory groups possible. Learning rate in the vertical component is comparable between groups as it a visual-only judgment for each. For the horizontal component, no comparison can be made between the training rate in the two groups because the DRT for the horizontal component judgment in the auditory group remained constant at near-ceiling levels. This was expected because this group had unambiguous, supra-threshold information about the horizontal component of motion direction from the purely horizontal auditory motion stimulus.

To assess overall change in thresholds as a result of training, we compared the change in DRT from pre-test to post-test performance in both components of motion direction, and tested for effects of training stimulus type (audiovisual and visual-only), in both trained and untrained locations. The pre- and post-tests used visual-only stimuli, so results in both components of motion direction are comparable between groups.

### 2.8.1. Statistics

Analysis was done by fitting a linear mixed effects model to the data and conducting post-hoc tests to determine the impact any factors had on the results. These models were fit using the lme4 package (Bates et al., 2014) in the R statistical software environment (version 4.0.5 (R Core Team, 2021)). The significance of each effect in these models indicates whether that factor had significantly different parameter estimates between its levels. For example, a linear mixed effects model was fit to the DRT in the training location and included pre- to post-test change, group assignment (audiovisual or visual-only) as well as component of DRT (horizontal or vertical), and interactions between them. If group assignment were to show as a significant factor, we could conclude that the group assignment had significant differences between factor levels. In this case, this would indicate a difference between audiovisual and visual-only.

In this model, since pre-test to post-test is coded as a factor and we primarily care about the change from pre-test to post-test, we can disregard any effects that do not involve an interaction with the the pre- to post-test term. If the main effect of the pre-test to post-test were significant, but the interaction between pre- to post-test and group assignment was not significant, we could conclude that while both group assignments show change from pre-test to post-test, there is no difference in the magnitude of this change between the groups.

In order to understand the effects further, post-hoc testing was done using the linear mixed effects model by calculating the estimated marginal means in R using the emmeans package (Lenth, 2021) and running interaction contrasts to determine whether the change from pre-test to post-test was different between groups, locations, and component of judgment. Detailed descriptions of the models used and contrasts calculated can be found in Section 3.

## 3. Results

### 3.1. Effect of training on the trained task

Training results from days 2-11 of the study were evaluated to assess the effect of the added auditory stimulus on learning rate. Because the audiovisual group was given supra-threshold information about the horizontal component of movement direction, horizontal judgments were near perfect, as was expected. For this reason, we focused analysis on the judgments of motion direction based upon the vertical motion component of the stimulus, which was statistically independent from the horizontal motion component. DRT for the vertical component was fit by a linear mixed effects model. The participant’s study ID was included as a random factor, which contributed to 76.6% of the overall variance in the model. The fixed effects in the model were group assignment, which was a factor with 2 levels, and day, which was a continuous numeric variable. The effect of day was significant (*p <* 0.001), indicating that performance increased overall during the training. Additionally, the interaction between day and group was significant (*p* = 0.017), meaning that the groups had significantly different rates of change. The marginal slope estimate for the audiovisual group was 3.513°/ day with standard error of 0.6042, and for the visual-only group the slope estimate was 5.646°/ day with standard error of 0.6409. The audiovisual group learned more slowly than the visual-only group for the vertical component judgment. A separate linear mixed effects model was fit for the training data within the visual-only group to compare slopes in the vertical and horizontal component. As in the previous model, DRT for both components was fit by the model, with the participant’s study ID as a random factor. This model owed 61% of its variance to random effects. The fixed factors include the two direction components of the judgment of motion direction (treated as independent factors, horizontal and vertical), and the day of training. This model showed a significant effect of day (*p <* 0.001) but no significant interaction between the component and days, indicating that the slopes for each component were not significantly different.

### 3.2. Transfer of learning to the visual-only condition

Days 1 and 12 of the experiment included a pre- and post-test in the training location without the presence of a cue. In the audiovisual group, learning achieved in the training task with the auditory cue failed to transfer to the post-test condition where the auditory cue was removed, as shown in Figure 5.

**Figure 5:**
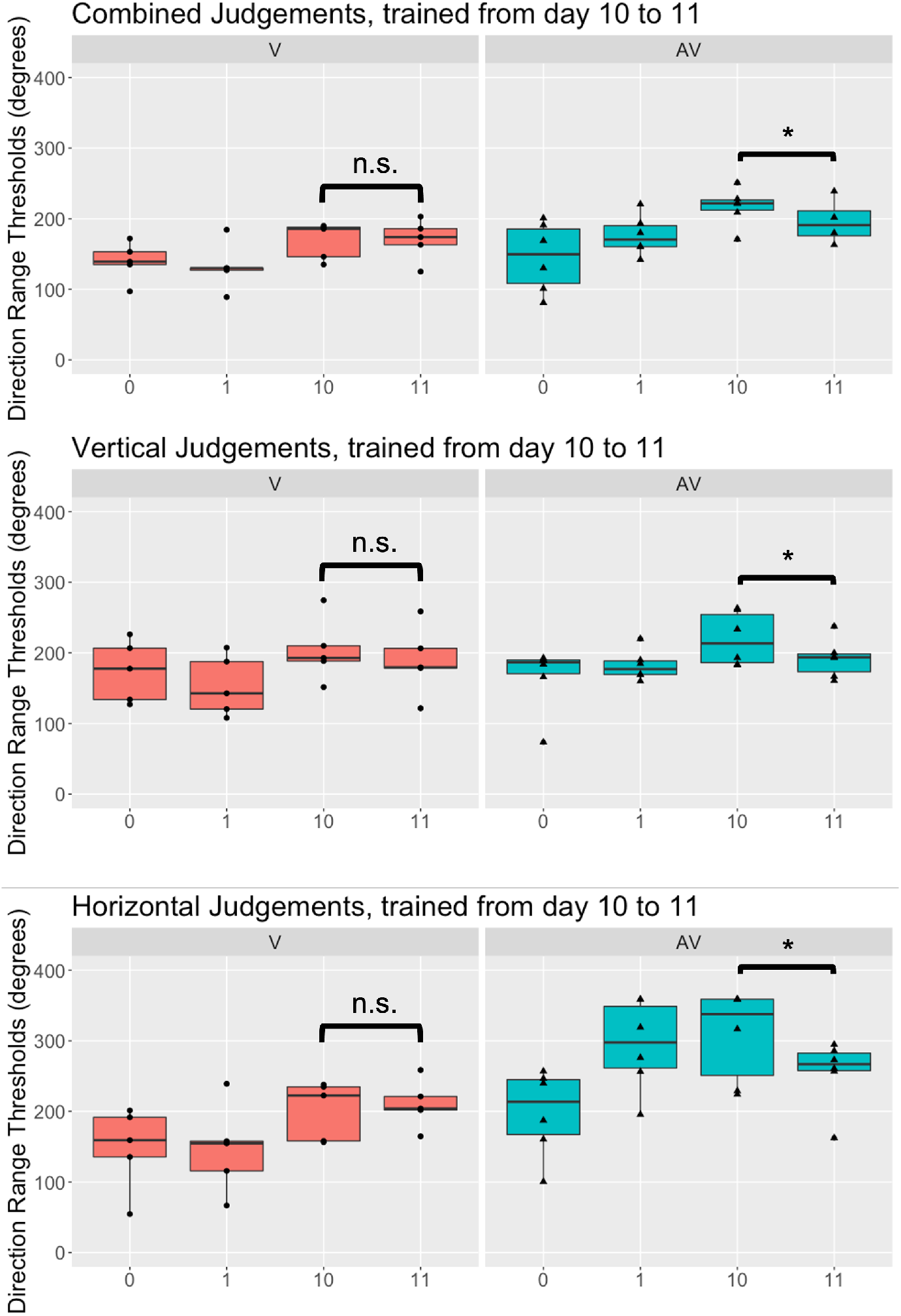
Task transfer in training location: Direction range thresholds for the over-all combined judgment of the motion direction, as well as the horizontal and vertical components of the motion direction judgment are averaged for each group (error bars are standard deviation, point markers represent individual participants). Direction range thresholds are shown on the first and final day of training (days 1 and 10), where the auditory cue was present for the audiovisual group, and in the pre-test and post-test, where the auditory cue was absent in both groups. The audiovisual group shows a significant difference (p = 0.02) between their performance on day 10 (cue present) and the post-test (cue absent) for all the components of the judgment.

To assess transfer of learning on the task including the auditory cue to the task without the auditory cue, a linear mixed effects model was fit to the change from the pre-test to the first day of training as well as the change from the last day of training to the post-test on the vertical component of the judgment. Participant’s study IDs were included as a random factor, and this random effect contributed 79% of the overall variance in the model.

The audiovisual group showed a significant change (p = 0.02) from day 10 of training, which included the auditory cue, to the post-test, which had no auditory cue. The audiovisual group had an estimated slope of -22 degrees of direction range, with standard error of 8 degrees of direction range, and the visual group had no significant change from day 10 to the post-test.

To examine transfer of learning to the task condition with no auditory cue in the horizontal component of motion, training data could not be directly compared to the post-test data for the audiovisual group. As the participants in the audiovisual group were provided an unambiguous cue to the motion direction during training, this group remained at ceiling for the duration of the training. Thus, their results on day 10 where they received the auditory cue are not indicative of their visual perception alone as the auditory cue provided full information about the horizontal component of the motion direction. However, by comparing the pre- and post-test results for the horizontal component, task transfer can be inferred. The pre- and post-test data for all participants, including both components of the motion direction judgment were fit using a linear mixed effects model. This model had the response variable of DRT, and included the individual participant ID as a random factor. The random effect contributed 31.3% of the overall variance in the model. The fixed effects were group assignment, location of test, and component of judgment as well as the interactions between them. In this model, no terms were significant, indicating that the effects of the levels of the included factors were minimal. To determine whether there was improvement within any of the treatment combinations, consecutive contrasts comparing the pre-test to the post-test were calculated for each group, location and judgment component combination.

In the trained location, both groups improved only within the horizontal component of judgment, as shown in Figure 6. The visual-only group showed a significant improvement (*p* = 0.030) in the horizontal judgment within the trained location. Change was estimated at 61.6 degrees of direction range with a standard error of 27.9. The audiovisual group also showed significant improvement (*p* = 0.028) in the trained location for the horizontal component, with the change estimated as 57.0 degrees of direction range, and a standard error of 25.5. In the training location, neither group showed significant improvement in the vertical component of the judgment as shown in Figure 6.

**Figure 6:**
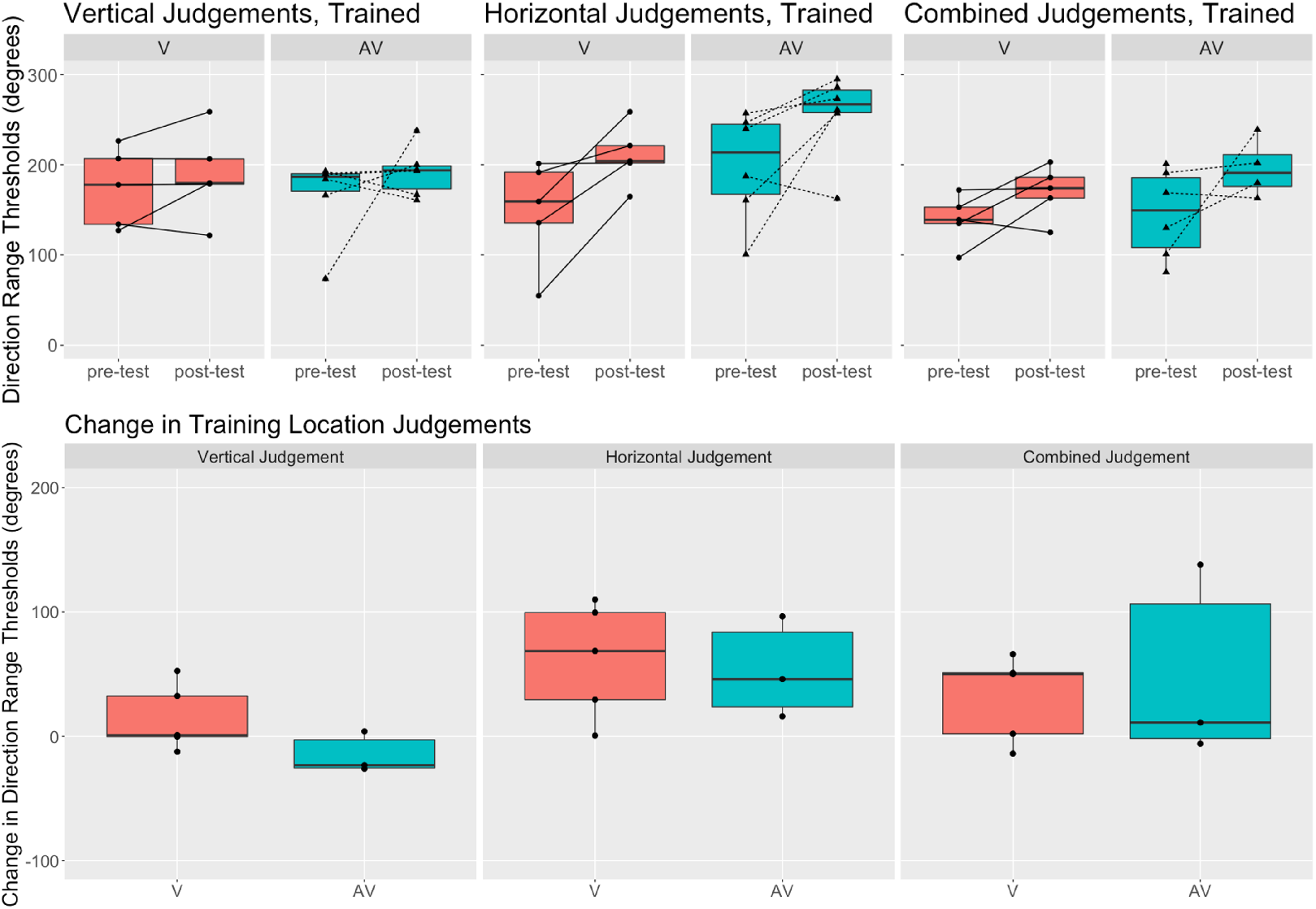
Pre-/post-test change in horizontal and vertical component judgments for both groups in the trained location. Individual participants are represented by round or triangle markers. The boxplots show range as vertical lines. The top row of plots show the direction range threshold performance in the pre-test and post-test for the vertical component, the horizontal component, and the combined components of the motion judgment, in both the visual-only group (left, orange) and audiovisual group (right, blue). The bottom panel shows the change in performance as post-test minus pre-test. Vertical judgments showed no significant change. Horizontal judgments in the visual-only group (orange) improved significantly with an estimated change of 61.6 ± 27.9 degrees (*p* = 0.03). The audiovisual group (blue) also improved significantly in the horizontal component, with an estimated improvement of 57.0 *±* 25.5 degrees (*p* = 0.028)

### 3.3. Transfer of learning to an untrained location

In the untrained location, only the audiovisual group showed significant change from pre-test to post-test, as seen in Figure 7. The audiovisual group showed significant improvement (p = 0.0145) in the horizontal component of judgment with change estimated at 52.0 degrees of direction range, and standard error of 20.8 degrees of direction range. The vertical component also showed near-significant improvement (p = 0.0514) for the audiovisual group with change estimated at 41.1 degrees of direction range with standard error of 20.8 degrees of direction range.

**Figure 7:**
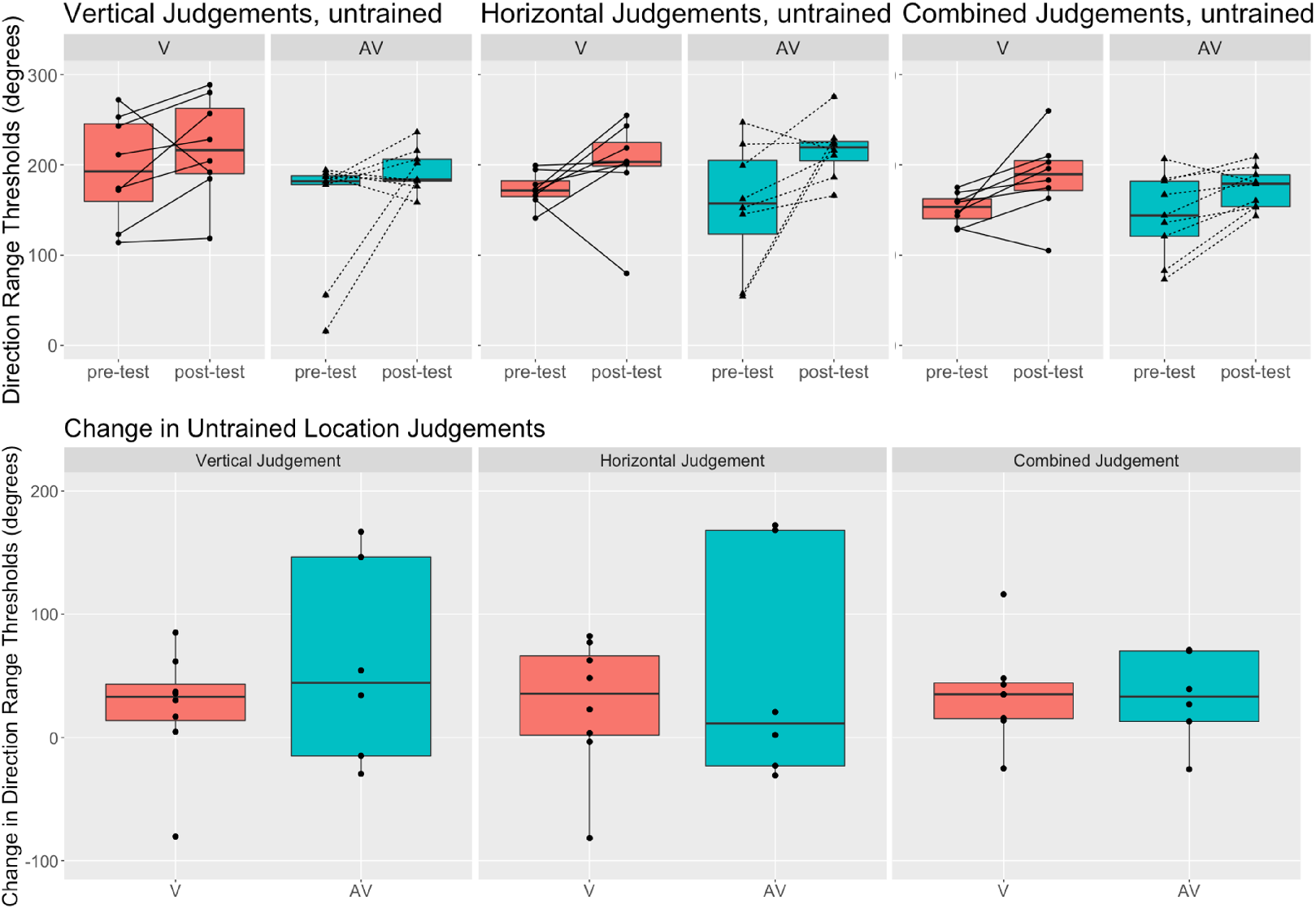
Pre-/post-test change in horizontal, vertical, and combined component judgments for both groups in the untrained location. Individual participants are represented by round or triangle markers. The boxplots show range as vertical lines. The top row of plots show the direction range threshold performance in the pre-test and post-test for the vertical component, the horizontal component, and the combined components of the motion direction judgment, in both the visual-only group (left, orange) and audiovisual group (right, blue). The bottom panel shows the change in performance as post-test minus pre-test. The visual-only group (orange) showed no significant improvement in either the horizontal or vertical component. The audiovisual group however showed significant improvement in the horizontal component estimated at 52.0 ± 20.8 degrees (p = 0.0145), and a near-significant change in the vertical component estimated at 41.1 *±* 20.8 degrees (p = 0.0514)

Despite the presence of main effects in the comparison of pre- and post-tests, when interaction contrasts are calculated to compare the *changes* between different factor levels, there is no significant difference between any of the groups, locations, or judgment components. For example, in the training location for the visual-only group, while the change from pre-test to post-test was significant for the vertical component of the judgment and not the horizontal component of the judgment, when comparing the estimates of these changes, there is no significant difference between them. This is likely due to the high noise level in the data compared to the effect size; and should be unsurprising given that the standard error in these estimates are around 50% of the estimated quantities.

## 4. Discussion

Results from this study show that visual training with an added auditory cue can still produce learning, but that learning may not transfer to performance in the absence of the cue. One notable exception is when learning did transfer to performance in the absence of the cue, it also showed transfer to an untrained location. This transfer to the untrained location was observed in the audiovisual group for the component of learning that showed task transfer.

Although audiovisual and visual-only groups both showed learning in the purely-visual, vertical component of the motion direction judgement, the audiovisual group demonstrated a notably slower learning rate in this task. It is possible that the observed difference in learning rates between the two groups is related to the allocation of attention during training. When training, the presentation of the vertical component was the same for both groups. However, the audiovisual group had additional information provided for the horizontal component in the form of the auditory cue. This added auditory cue may have been detrimental to learning. Although attention can be a strong driver of visual perceptual learning (Gutnisky et al., 2009; Nguyen et al., 2020), it is also a limited cognitive resource, and the demands that audiovisual integration places on attention are especially strong when the auditory and visual stimuli are spatially and temporally coherent (Van der Burg et al., 2008; Driver and Noesselt, 2008; Fleming et al., 2020). Moreover, information about the horizontal component of motion was irrelevant to correct judgement of the vertical component, which is the dimension that drove task difficulty. These observations suggest that the cross-modal spreading of attention may have led to an irrelevant auditory stimulus capturing more of the participant’s attention away from a visual task (Zimmer et al., 2010a,b).

Following training, the audiovisual group showed transfer of learning to the untrained location, albeit only for the horizontal component of motion direction judgments. This observation is based on a comparison between the pre-test performance and the post-test performance for the reason that, in the presence of the supra-threshold auditory cue to horizontal motion direction, the group’s judgments of horizontal motion direction were at ceiling performance during training and were therefore uninformative about transfer. It is notable that, although the auditory cue to horizontal motion direction was present during training, it was not present during the pre and post-tests. Nevertheless, the transfer of learning occurred. This offers one explanation for why improved judgements of vertical motion direction did not transfer from training to the post-test. Attention can be enhanced by congruent cross-modal stimuli (Ee et al., 2009), and in our task the horizontal component of the visual motion was congruent to the auditory motion while the vertical was not. Engagement of endogenous attention can improve transfer of visual perceptual learning (Nguyen et al., 2020), and it is possible that this increased attention to the horizontal component due to the cross-modal cue led to a transfer of learning to the task condition where no cue was present. In summary, the transfer of only the horizontal component of judgments is consistent with our initial hypothesis that audiovisual integration can support better transfer to untrained locations, perhaps by engaging the top-down mechanisms involved in multisensory integration(Powers III et al., 2016b; Seitz et al., 2006).

The data reveal a similar outcome for transfer to the trained location; comparison of pre and post tests at the trained location show improved judgments of horizontal motion direction, but not vertical, despite the observed improvements for both during training, for both groups. One possible explanation is that experimental design required the feedback structure of the pre/post training paradigm to be slightly modified from that used during training. In all conditions, feedback was provided at the end of each trial both through an indicator sound which was positive for a correct vertical judgement and negative for an incorrect vertical judgement, and by a visual representation of the correct vertical and horizontal motion directions as an arrow pointing to the correct quadrant (NE, SE, SW, NW; see Fig. 1). During training, feedback had an intuitive relationship to task difficulty, which was modulated in accordance with a single staircase. In contrast, two interleaved staircases independently measured the vertical and horizontal components of motion sensitivity. As a consequence of, the relationship between feedback at the end of each trial and task difficulty on the next trial was unreliable, and this may have contributed to a loss of a sense of progress. Negative feedback, even when false, has been shown to negatively affect learning performance (Amitay et al., 2015), and if the slight change to the feedback was perceived as subtly negative it may have disrupted gains made in training.

Among the many potential differences in methodology that might explain why this study only partially replicated the findings of Seitz et al. (2006), which also investigated the role of an auditory component on visual learning, we speculate that the most influential difference may have been that of task difficulty. Task difficulty has been shown to have a strong effect on the quality of perceptual learning (Ahissar and Hochstein, 1997). The auditory facilitation of visual perceptual learning was demonstrated in Seitz et al. (2006) using a task that remained at a fixed difficulty level, rather than a level adaptively determined through a staircase, as in the present study. The subsequent data suggest that the task was quite difficult even at its lowest noise settings. Moreover, in recruiting for the study, almost a third of participants could not complete the titration task described in Section 2.6.1 and were excluded from the study. In similar tasks completed during pilot testing which only used a left/right discrimination versus a four-way discrimination task, dropout levels were much lower, and almost exclusively due to self-exclusion due to discomfort in the VR headset. The high difficulty level was set deliberately in part to encourage audiovisual interactions which have been shown to be strengthened when the visual information is of lower quality (De Niear et al., 2016; Parise and Ernst, 2017; Deneve and Pouget, 2004).

The difference in task difficulties may also explain why participants in this study also demonstrated notably different learning curves from those in Seitz et al. (2006). This is true even though Seitz et al. (2006) experiment also involved a 10 day training period for motion direction discrimination, and even though the resulting overall attainment between the visual and audiovisual groups was similar to the levels found in this study. Moreover, although the audio / audio visual training groups in this study did not demonstrate different learning curves, the audio / audio visual training groups in Seitz et al. (2006) did. One possibility is that the use of an adaptive staircase provided a more uniform level of difficulty across the two groups, while the use of a fixed-difficulty design meant that the addition of the auditory component made the audiovisual group’s task relatively easier than the task presented to the visual-only group. The difference may also be attributed to the current tasks use of a 4AFC task paradigm, which differs from the 2AFC task used in Seitz et al. (2006). While most literature around task difficulty in perceptual learning centers around specificity of training, the results from (Wang et al., 2013) show shallower training slopes for a higher precision task than for the same task with lower precision. As task precision required for a 4AFC task is higher than for a 2AFC task, a longer training period may be needed to produce more consistency between participants in this 4AFC task.

In summary, this study found that the benefit from the addition of an auditory component to visual perceptual learning was mediated by several factors, including the spread of attention across the modalities, the quality of feedback, and task difficulty. This suggests that sound is not automatically a facilitator of visual perceptual learning, but rather that multisensory engagement is one factor among many which can impact perceptual learning, and that all these factors must be carefully tuned if the goal is either to optimize the rate of learning or to optimize the transfer of learning to untrained portions of the visual field.

